# The emergence of medusa-specific cell states in the scyphozoan *Aurelia coerulea*

**DOI:** 10.1101/2023.08.24.554571

**Authors:** Oliver Link, Stefan M. Jahnel, Kristin Janicek, Daniel Guerguerian, Johanna Kraus, Juan D. Montenegro, Bob Zimmerman, Brittney Wick, Konstantin Khalturin, Alison G. Cole, Ulrich Technau

## Abstract

The life cycle of most medusozoan cnidarians is marked by the metagenesis from the asexually reproducing sessile polyp and the sexually reproducing motile medusa. At present it is unknown to what extent this drastic morphological transformation is accompanied by molecular changes in the cell type composition. Here, we provide a single cell transcriptome atlas of the cosmopolitan scyphozoan *Aurelia coerulea* focusing on changes in individual cell states during the transition from polyp to medusa. Notably, this transition is marked by an increase in cell type diversity, including an expansion of neural subtypes and the appearance of striated muscles. We find that two families of neuronal lineages are specified by homologous transcription factors in the sea anemone *Nematostella vectensis* and *Aurelia coerulea*, suggesting an origin in the common ancestor of medusozoans and anthozoans about 500 Myr ago. Our analysis suggests that gene duplications might be drivers for the increase of cellular complexity during the evolution of cnidarian neuroglandular lineages and highlight the close relationship of neurons and muscles. One key medusozoan-specific cell type is the striated muscle in the subumbrella. Evaluating muscle types by fibre anatomy and gene expression validation of their individual molecular profiles made it possible for the first time to investigate transcriptome differences between smooth and striated muscles. Although smooth and striated muscles are phenotypically different, both have a similar regulation of the contractile complex, reminiscent to the regulation of smooth muscles in bilaterians. This is in contrast to bilaterian striated muscles, where the regulation of muscle contraction involves Ca**^2+^** binding troponins and their interaction with Tropomyosin. This suggests that smooth muscle contraction regulation is ancestral and the use of troponins in striated muscles only evolved in bilaterians.

**Teaser:** Single cell transcriptome atlas across the jellyfish life cycle reveals emergence of novel medusa-specific cell types associated with expression of medusa-specific paralogs.

## Introduction

Cell type diversification is a hallmark in the evolution of metazoans, and the principles that underlie these processes are still barely understood. Advances in single cell sequencing allow the cataloguing of differential transcriptome usage across cell types from entire organisms (reviewed in (1,2)). To understand the evolution of cell type diversification that coincided with the emergence of complex body plans in the animal kingdom, members of basal branching animal phylum Cnidaria hold the key to reconstructing ancestral cell states present during the explosion of animal morphotypes that remain in modern day. Cnidarians are the phylogenetic sister group to Bilateria (3–5), which comprise most of all animal phyla. In this regard understanding the phylum Cnidaria is required for the identification of key bilaterian traits.

The phylum Cnidaria is composed of the subphylum Anthozoa, which encompasses sea anemones, corals and sea pens and its sister group, the Medusozoa. Except for Staurozoa (6), Medusozoa differ from the anthozoans by typically having a triphasic life cycle that includes a free-swimming larva, an asexually reproducing polyp, and a free-swimming sexually reproducing jellyfish (a.k.a. medusa) stage (6,7). The latter is not present in anthozoan clades. The formation of these two distinct adult life forms in anthozoans and medusozoans has fascinated researchers for more than a century (6,8–10). The life cycle of the anthozoan *Nematostella vectensis* has been characterised at a single cell transcriptome level by recent studies (11–13). However, most medusozoan life cycles are characterized by a drastic metagenesis from the polyp into free-swimming jellyfish which provides opportunities to investigate the evolutionary principles underlying the development of any new cell types that emerge during the switch from a sedentary to a free-living lifestyle.

Within Medusozoa, the hydrozoans, scyphozoans and cubozoans form medusae in quite diverse ways, raising the question whether they are homologous or convergent (8,14). Recent single cell transcriptome studies in hydrozoans have analysed the freshwater polyp *Hydra* (15), which has lost the medusa stage, the medusa stage of *Clytia hemisphaerica* (16), and a series of medusa, ‘four leaf structures’, ‘cysts’ and polyps characterizing reverse development in *Turritopsis rubra* (17). The latter study also compares *T. rubra* with the medusa of *Aurelia coerulea* collected from the waters near Yantai China. The molecular changes underlying the transition from polyp to medusa have been characterised via bulk transcriptome sequencing and show that each life stage is characterized by the expression profile of different transcription factors that highlight individual molecular pathways (18–20). However, changes at the cell type level during the metagenic transition were not investigated in these studies. Here we present a single cell characterisation of the metagenesis in a scyphozoan, the moon jelly *Aurelia coerulea* (formerly sp.1), from a strain of unknown origins that has been kept in the laboratory for over 25 years.

In most scyphozoans like *Aurelia,* metagenesis occurs via polydisc strobilation, an asexual horizontal fission process along the oral-aboral axis of the polyp which enables the organism to produce up to 20 free-swimming individuals from a single polyp (21). Striated muscle and a set of neurons and gravity sensing cells are considered to be medusa-specific and have been studied from the perspective of morphology (22–26), physiological features (27,28), and specific gene expression profiles (29,30). Scyphozoan jellyfish nervous systems consist of two individual nerve nets termed the “motor nerve net” (MNN) and the “diffuse nerve net” (DNN) (27,31). Nerve nets exclusively use chemical synapses (32) and are “through conducting” (33), although less is known about the DNN than the MNN. The MNN shows a non-polarized transmission of excitation with bi- or multipolar neurons (27,31). Together, both nerve nets are responsible for either fast (MNN) or slow (DNN) contraction waves of the jellyfish umbrella (34) and regulate muscle contraction of circular umbrella muscles by the MNN or marginal radial muscles by the DNN. While it has been shown that FMRFamide antibodies specifically label cells of the DNN and non-modified / modified alpha or beta-Tubulin chains label cells of the MNN (23,28,35), the number and character of the neuron types that compose these circuits is unknown.

In addition to the D/MNN nerve nets, the medusa nervous system includes also the sensory rhopalia. The rhopalia connect with both DNN and MNN neurons that innervate the striated musculature in the epidermis of the medusa umbrella and act as a pacemaker enabling jellyfish navigation and swimming behaviour (22,28,31). This rhopalia nervous system develops during polyp-medusa metamorphosis and is composed of specialized light-(pigment cup) and gravity-sensing (lithocyte/statocyst) cells, segregated into individual compartments with different developmental origins (22). Rhopalia development involves the gene expression of *otx1*, *pit1* and *brn3* in the pigment-cup (29), three transcription factors with a conserved role in brain development in bilaterians. Effectors downstream of the sensory system are the muscle cells. Striated muscles show a distinct sarcomere organization which in *Aurelia* become ultrastructurally evident only upon the ephyra stage (7,26), with minor exceptions (24). Additional cell types include the digestive cells housed within the gastric filaments of the *Aurelia* ephyra and medusa stage, which are identifiable from the expression of digestive enzymes such as *chitinase*, *protease* and *pancreatic lipases* (36). Lastly, six different types of cnidocytes have been recorded in the species *Aurelia aurita* (37), but it is not clear whether all six cnidocytes are present in both polyps and medusa. These have been categorized as a-isorhiza, A-isorhiza, O-isorhiza, eurytele, polyspiras and birhopaloid (37), the most common being a-isorhizas and euryteles.

In this study, we provide a molecular foundation for investigating mechanisms of cell type diversification in Scyphozoa by generating a cell type atlas of the scyphozoan model, *Aurelia coerulea* (formerly sp.1). We used single-cell RNA sequencing to produce whole animal, as well as tissue specific single-cell RNAseq libraries that cover the transition from polyp (also called scyphistoma) to jellyfish. We characterise the composition of identified cell type families across the life cycle and identify cell types that define the jellyfish as free-swimming individuals, including various neuronal subpopulations and striated muscles, confirming anatomical reports, providing here complete gene expression profiles of the different neuronal cell populations. We found parallels with anthozoans in terms of reported cellular complexity with regards to neural composition and cnidocyte specification pathways. And finally, we characterise the molecular signatures of the medusa striated muscles compared to the smooth muscles of the polyp and identified a set of genes that is shared between both muscle phenotypes. These genes correspond to a shared contractile complex between both types. With this analysis we set the foundation for comparative studies addressing evolutionary origins of medusa specific cell types or investigating the assembly of the striated muscle phenotype in Cnidaria.

## Results

The typical life cycle of medusozoans involves the transition from a sessile polyp (scyphistoma) to the sexually active jellyfish (medusa). This not only is a dramatic reorganisation of the body plan, but it is also accompanied by the emergence of medusa-specific cell types and potentially the disappearance of polyp-specific cell types. We generated sixteen single cell RNA seq libraries using the 10x Genomics platform: twelve from different stages of the life cycle of *Aurelia coerulea* and four from different tissues of the medusa. We note that the first morphological sign that a polyp is competent to strobilate is the expansion of the body column into four pouches and resembles a cloverleaf from above; we refer to this stage as the “clover-polyp”. Our single cell dataset includes two replicates of the sessile polyp, a clover-polyp, two replicates of clover-polyp with tentacle bulbs separated from the body column and processed independently, an early and late strobilating polyp, newly liberated ephyra juvenile, fed ephyra, and the newly metamorphosed medusa (1 month, ∼1.5cm diameter); four tissue libraries from a growing medusa (∼8cm diameter) including the umbrella (excluding the margin), the margin (including tentacles), the manubrium (including the arms and mouth), and the gastrodermis (including future gonad) (**Fig. 1A**). Despite its larger size, this animal was still reproductively immature and so no gonadal tissues were collected. All processed cell suspensions showed a viability estimate of over 75% using a fluorescein/propidium iodide assay for quantification. We mapped the sequence data to the *Aurelia coerulea* isolate AC-2021 genome (NCBI: GCA_039566865.1) annotated with Augustus. Each single cell library was sequenced to at least 70% saturation, from which we detect 24,028 / 25,954 gene models of which 20,432 are represented by 10 or more unique reads across the entire dataset. With these data we provide an inventory of cellular changes that occur during metagenesis.

**Fig. 1.**
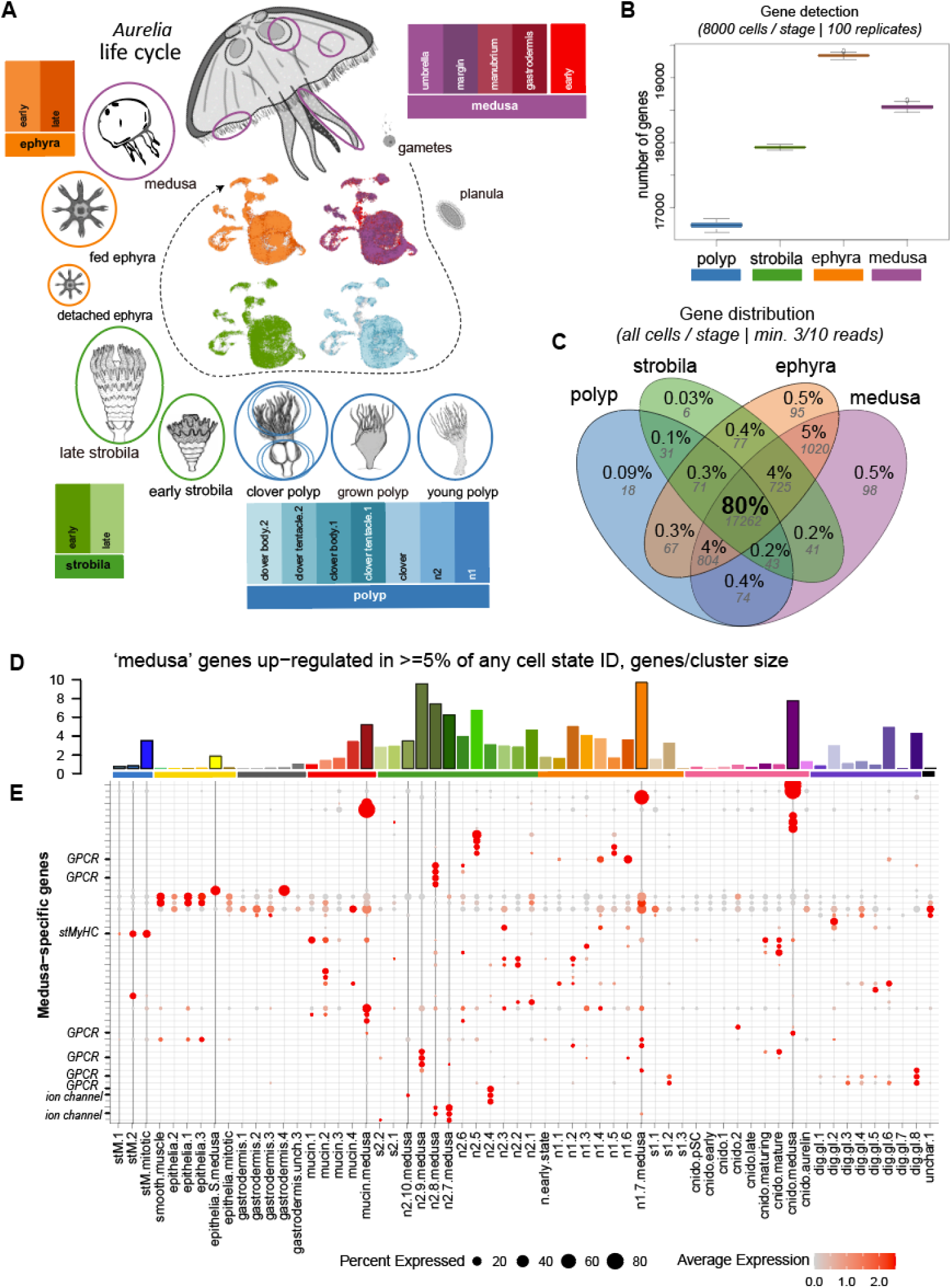
Single cell transcriptome atlas covering the polyp-medusa life cycle transition demonstrates shifts in genome usage between key life history stages. (**A**) Twelve single cell transcriptomic libraries were generated from cell dissociations of eight sequential life history stages and four tissue compartments from a sub-adult medusa. Samples are color-coded according to their life-cycle. In the center UMAP cell plots of the full dataset separated by life cycle illustrates the distribution of cells from each library (**B**) Distribution of genome use across the life cycle. Number of gene models (>3/10 reads) detected for cells binned according to stage, downsampled to 8000 cells representing each stage, with 100 replicates. (**C**) Venn Diagram illustrates the overlap of detected genes from a single replicate of (B). (**D**) The number of medusa-specific genes expressed in each cell state, normalized by cluster size. Cell partition and cell states are colored as in Figure 2. **E**) Expression of genes not detected in either polyp nor scyphozoa in figure (C), filtered to include only genes that are up regulated in at least 5 % of any cell state in the dataset. Medusa-specific cell states are highlighted with black outline.

### Transition to free-swimming medusa is accompanied by increased cell type complement

We first asked whether we could identify gene sets specific for each life-cycle stage (polyp: 16644 cells; strobila: 9332 cells; ephyra: 15299 cells; medusa: 21955 cells) by comparing stage-specific samples against one another. To accomplish this, we randomly down-sampled to 8000 cells (with replacement) per stage from the collection of stage-specific samples and then calculated the number of genes detected at each stage. We confirmed this observation with 100 replicates; the number of expressed genes detected was consistently ∼10% lower in polyp samples with respect to the other life cycle stages (**Fig. 1B**). We note that all samples derived from polyp cells overall express fewer genes compared to other stages despite similar sequencing depth (**Fig. S1A,B**). We examined the overlap of the detected gene sets and found ∼6% of expressed genes are not detected in the polyp nor strobila samples (**Fig. 1C**), indicating that a small portion of the coding genome may be activated only in the medusa phase of the life cycle. We queried the identity of this medusa-specific gene set and found an enrichment in signalling and ion channels (**Fig. S1C, Data S1.1**), suggesting a possible increase in neuronal diversity within the medusa compared to the sessile polyp stage. The vast majority of these ‘medusa’ genes are expressed at very low levels (< 50 reads total) and thus could easily represent false negatives in the polyp datasets, and are detected across all cell type identities. We filtered this gene set to include only those up-regulated in at least 5% of any cell state identity (as described below), and examined their distribution normalized by cluster size (**Fig. 1D, Fig. S1C**). The resulting 49 genes are detected in cell-state specific expression profiles, with an over-representation in medusa-specific cell states (**Fig. 1D,E**). This gene set includes 5 rhodopsin-like GPCR genes and a striated-type myosin heavy chain (*stMyHC-5*). We also examined the set of differentially expressed genes across all four life cycle stages (**Fig. S2, Data S1.2, S1.2b**) and found that the polyp-specific gene complement is enriched for prostaglandin biosynthesis (**Fig. S2B**), suggesting that this hormonal pathway might be involved in the maintenance of the polyp state. Consistent with this observation, it has been demonstrated that blocking prostaglandin signalling with indomethacin can induce strobilation in this species (38). The medusa stage, with its large muscular swimming bell, is characterised by an enrichment of collagen and muscle-related genes (**Data S1.2**). The upregulation of collagen genes reflects the thickening of mesoglea in the medusa, while the muscle-related genes are employed in the expansion of the muscle field in the subumbrella of the medusa.

We next sought to generate an inventory of the cell diversity present at each point of the life cycle. All sixteen datasets were integrated using harmony PCA alignment (**Fig. 1A, S3**). We identified 9 broadly defined cell populations, for which we assign identities by extrapolating from up-regulated gene lists (**Data S1.3, S1.3b**). Principal cell populations include cnidocytes (pink), neurons (green), digestive (purple) and mucin (red) glands, striated muscle (dark blue), and two large populations that represent the inner (grey: gastrodermis) and outer (yellow: epithelia) cell layers, and one currently uncharacterized cell population (black: unchar.1) (**Fig. 2A**). The inner epithelia, or gastrodermis, expresses several collagens that is a characteristic of the inner cell layer of anthozoans (39); the outer cell layer houses the ring musculature and is rich in contractile proteins. The striated muscle cluster is also rich in contractile protein and is the only principal cell population absent from the polyp-derived samples (**Fig. 2C**). The mucin gland expresses mucin-like-proteins, whereas the digestive gland expresses other digestive enzymes, and the neural cluster expresses *synapsin* and other conserved known neural regulators such as *ashA*. The cnidocytes express mini-collagens and are enriched in pathways targeting the endoplasmic reticulum (40). To obtain a deeper characterization of cell type dynamics in this system, we investigated the substructure of each population in terms of cell type distribution and up-regulated genes and DNA binding molecules (**Data S1.4:10**). We consider all distinct transcriptomic clusters identified here as representative of cell type states, as in some cases identified clusters could represent different cell states of the same cell type. We used the aligned principal components from the integrated dataset to build a cell state tree for all identified clusters (**Fig. 2B**). The cell state tree indicates the majority of transcriptomic identities cluster according to partition and thus can be considered cell type families. In total we find 12 cell type states that are not represented (<3 cells) in the polyp scRNA sequencing libraries. We refer to these as medusa-specific cell states, all of which first appear within the strobila tissues and include three states of striated muscle, four are neural sub-types and one is a neural-related secretory cell type, and there is a single medusa-specific state in the epithelia, mucin, digestive gland, and cnidocyte partitions (arrowheads; **Fig. 2B**).

**Fig. 2.**
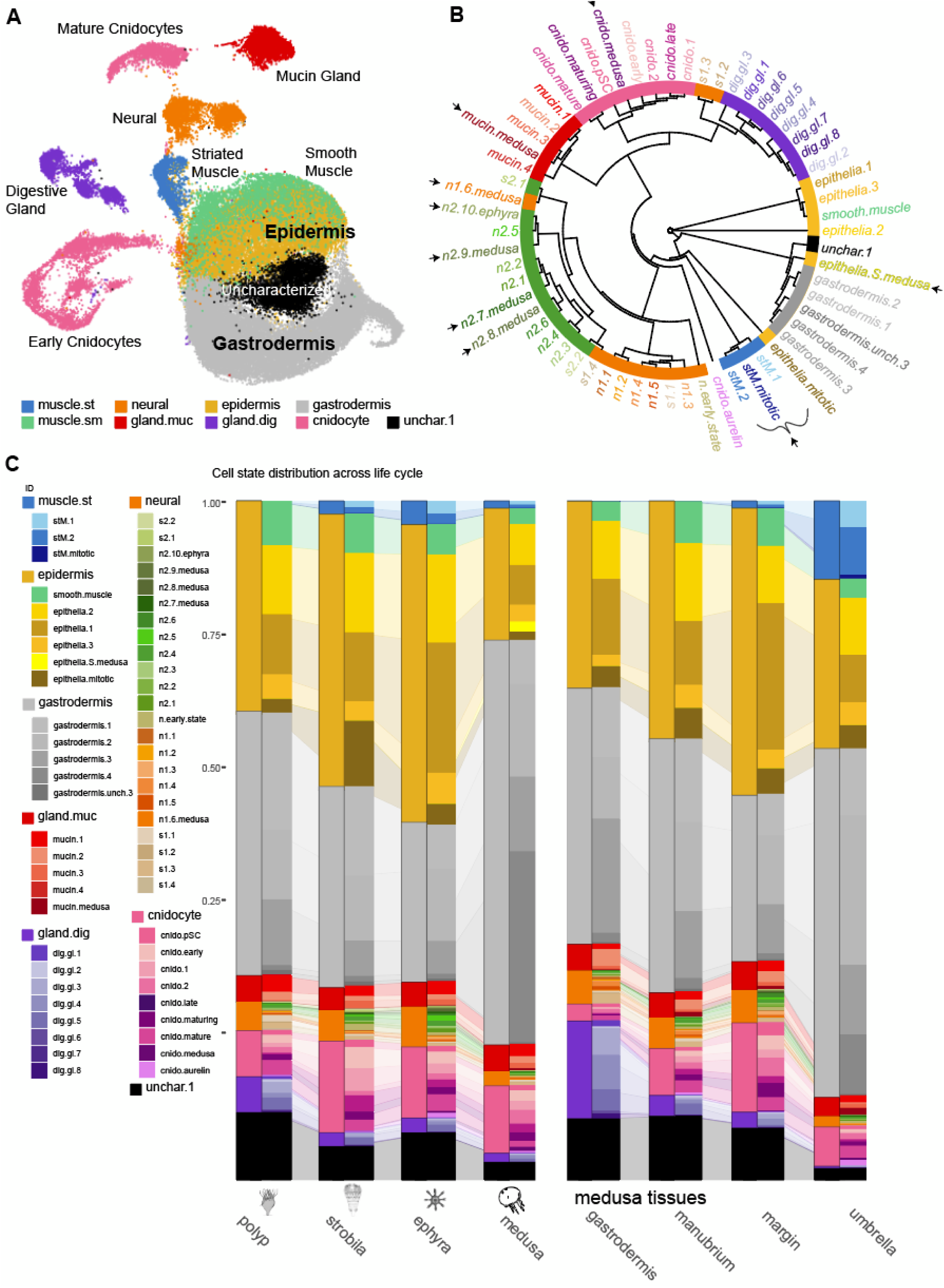
Integrated cell atlas permits identification of cell states across the life cycle. (**A**) UMAP representation of principal cell type partitions in the dataset. (**B**) Cell state tree illustrating the principal cell type families present in the dataset. Medusa-specific cell states are highlighted with arrowheads. (**C**) Distribution of cell type states across all life cycle stages plotted as a fraction of all cells captured. Both cell type family partitions and identified cell type states are shown.

Overall, the relative proportion of cells identified as the inner and outer epithelia remains constant across the life cycle. The polyp stage is characterised by having a larger proportion of digestive gland cell types, which then become localised to the gastric filaments, manubrial arms, or mantle margin in the medusa (**Fig. 2C** shades of dark purple; **Fig. S4A,B**). One digestive cell type is largely restricted to the polyp samples (*dig.gl.7*: yellow **Fig. S4C**), while there are no digestive cells in the umbrella sample (**Fig. S4C**). Digestive gland cell types, except *dig.gl.8*, share the expression of an *achaete-scuteC* (*ashC)* ortholog (**Fig. S4D**). Each sub-type has a different protease profile (**Fig. S4E; Data S1.10**), for example *dig.gl.3* expresses *chitinase-1* and are localized to the digestive filaments (**Fig. S4F**). Mucin subtypes (**Fig1G:** reds) specifically express a mucin-like protein (*mlp-like*) and *neuroD*-ortholog (**Fig. S5**), which is characteristic for neuronal cell populations in other systems (11,41–44). The mucin-producing population also contains one medusa-specific subtype (**Fig. S5**). Additional medusa-specific cell types include the striated muscle and several cnidocyte and neuronal subtypes described below.

### Cnidocytes highlight conserved features of cnidogenesis between anthozoans and scyphozoans

Recently we reported that within the sea anemone *Nematostella vectensis*, specification of the distinct cnidocyte types is marked by a diverging transcriptomic profile corresponding to the formation of the different capsule types, which then undergo a molecular switch demarcated by up-regulation of *GFI1B* and converge upon a secondary neural-like expression profile (11). Notably, we find a similar forked trajectory within the cnidocyte population of *Aurelia*. (**Fig. 3A**). A cluster of SoxC expressing ‘early’ cells separate along two principal trajectories (*cnido.1*, *cnido.2*), which then converge upon a second mature transcriptomic phenotype upon activation of jun/fos (**Fig. 3E**). We combined the *Aurelia* cnidocyte subset with that from the updated cell atlas for *Nematostella* (12), using the set of identified 1:1 orthologs as determined by OMA (45). We first removed cell clusters annotated as being ‘planula’ or ‘gastrula’ from the *Nematostella* dataset (**Fig. 3D**). We then aligned the principal components across all samples from both species using the harmony algorithm (46) and used these components to generate a cell state tree (**Fig. 3C**). Both early and late phase cnidocyte cell states from both species cluster together, whereas amongst the sub-type specification pathways we find support for *Aurelia* sub-type *cnido.1* and *Nematostella* sub-type *nem.1* with moderate support. The remaining *Nematostella* sub-types cluster together to the exclusion of the *Aurelia* subtypes. These data suggest that *Nv.nem1* and *Ac.cnido1* represent the isorhiza, which are proposed to be the only cnidocyte type shared between anthozoans and medusozoans (47). We identify a temporal sequence of transcription factor expression that follows the specification trajectories of all cnidocyte subtypes (**Fig. 3E**). We find the earliest cell states express two myc orthologs (*myc1* and *myc4*), a high mobility group (HMG) protein (*soxC*), a prdm13 ortholog (*prdm13a*), and a C2H2-type zinc finger protein (*ZN431-like-1*), similar to *Nematostella* (**Fig. 3E**). The end of the separate specification trajectories in both species is marked by *paxA* expression and activation of JUN/FOS orthologs, followed by expression of a *gfi1b* zinc finger ortholog marking the switch to a more neural-like transcriptomic profile and activation of *soxA* and additional *myc* orthologs (**Fig. 3F**). One core difference between the species is the late activation of *myc3* in *Aurelia* compared to *Nematostella*, whereas all other *myc* orthologs have similar temporal expression profiles within the cnidocyte lineages (**Fig. 3E, S6**). This indicates that the overall genetic trajectory of cnidocyte development is conserved between *Nematostella* and *Aurelia*, despite marked differences in their subtypes and their morphology.

**Fig. 3.**
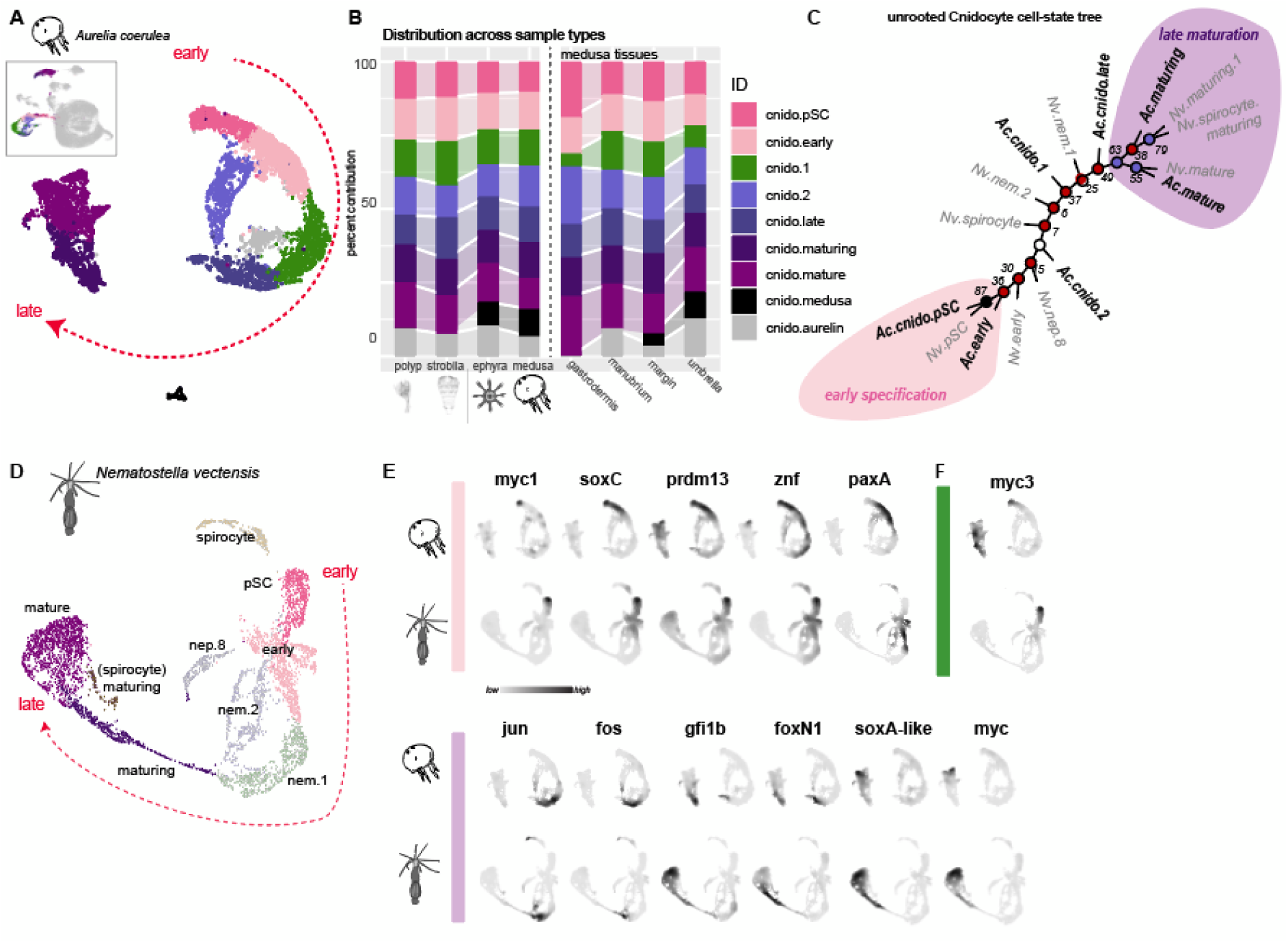
Cnidocyte specification pathways indicate two major capsule types. (**A**) Cnidocyte trajectory (UMAP) and (**B**) distribution (bar chart). Inset: location of subset within the full dataset. (**C**) Neighbour-joining cell state tree of cnidocyte profiles from *Aurelia* and *Nematostella* further indicate putative homologies between early (pink highlight) and late (purple highlight) phases of cnidogenesis. (**D**) Cnidocyte complement in *Nematostella vectensis* adapted from (14). The planula-specific cell clusters are removed. (**E**) Expression profiles of core transcription factor orthologs in *Aurelia* (top) and *Nematostella* (bottom) indicative of both early specification and late maturation. Paralogs show similar expression patterns in both species except for myc3 (green in (**F**)), which is expressed early in *Nematostella* and late in *Aurelia*. Bootstrap support values in (C) are indicated at the nodes; black indicates >80%, purple <80>50%, and red nodes are supported by <50% bootstrap.

### *Aurelia* neural complement reveals two neuro-secretory classes with similarities to anthozoan neuroglandular populations

The anthozoan *Nematostella vectensis* has two principal neural sub-families that have been described that correspond to those with *insulinoma* expression (n1) and those with *pou4* expression (n2) (11,12,48). The *Aurelia* single cell data contains a partition with neural properties that can be similarly separated into distinct transcriptomic profiles classified here into class 1 and class 2 subtypes (**Fig. 4A,B**). Here we define transcriptomic profiles as putatively ‘neural’ (“n”) based on the expression of voltage-dependent ion channels, whereas all other profiles are considered non-neural secretory “s” cell types (**Fig. 4C**). GO-terms associated with up-regulated gene sets specific to each sub-type largely supports this distinction and further suggest the “s” cells to be post-synaptically connected to the “n” states (**Fig. S7**). Our cell state tree suggests that all class 2 neurons form a united cell type family, whereas the class 1 neurons show closer affinities with other secretory cells (**Fig. 4D**). Similar to the distribution described in *N. vectensis*, *A. coerulea* class 1 neurons (orange) are *insulinoma* (*ins1*) positive and class 2 neurons (green) are *pou4* positive (**Fig. 4F**). Class n1 includes 6 putative neuronal sub-types: “*n1.1*” and “*n1.2*” are enriched in LWamide expression, “*n1.3*” and “*n1.6.m*” express putative acetylcholine receptors, “*n1.4*” expresses a putative netrin receptor A homolog, and “*n1.5*” expresses transcripts indicative of ciliation (**Data S1.9, Fig. S7**). The class n1 family also includes non-neural secretory cell types (“*s*”), which are enriched in genes associated with digestion and extracellular matrix production (**Data S1.9**). These data suggest a close relationship between neurons and gland cells, like what has been suggested in other cnidarians (11,15). In *Aurelia*, most neurons in the polyp stage are class 1 (**Fig. 4E**: shades of orange - ∼25% of all neural cells), with over 50% of polyp cells in the neural partition being of the s1 secretory variety (**Fig 4E**: shades of brown). These putative secretory cells are found primarily in the gastrodermis and manubrium samples of the medusa. Class 1 neurons in the medusa are also most prevalent within the gastrodermis. Thus, like that described for the anthozoan *Nematostella vectensis* (11,12), Class 1 neurons and related secretory cells comprise the predominant type of neuroglandular cells in the polyp stage. Furthermore, these are the primary neuroglandular cells within the gastrodermis of the medusa.

**Fig. 4.**
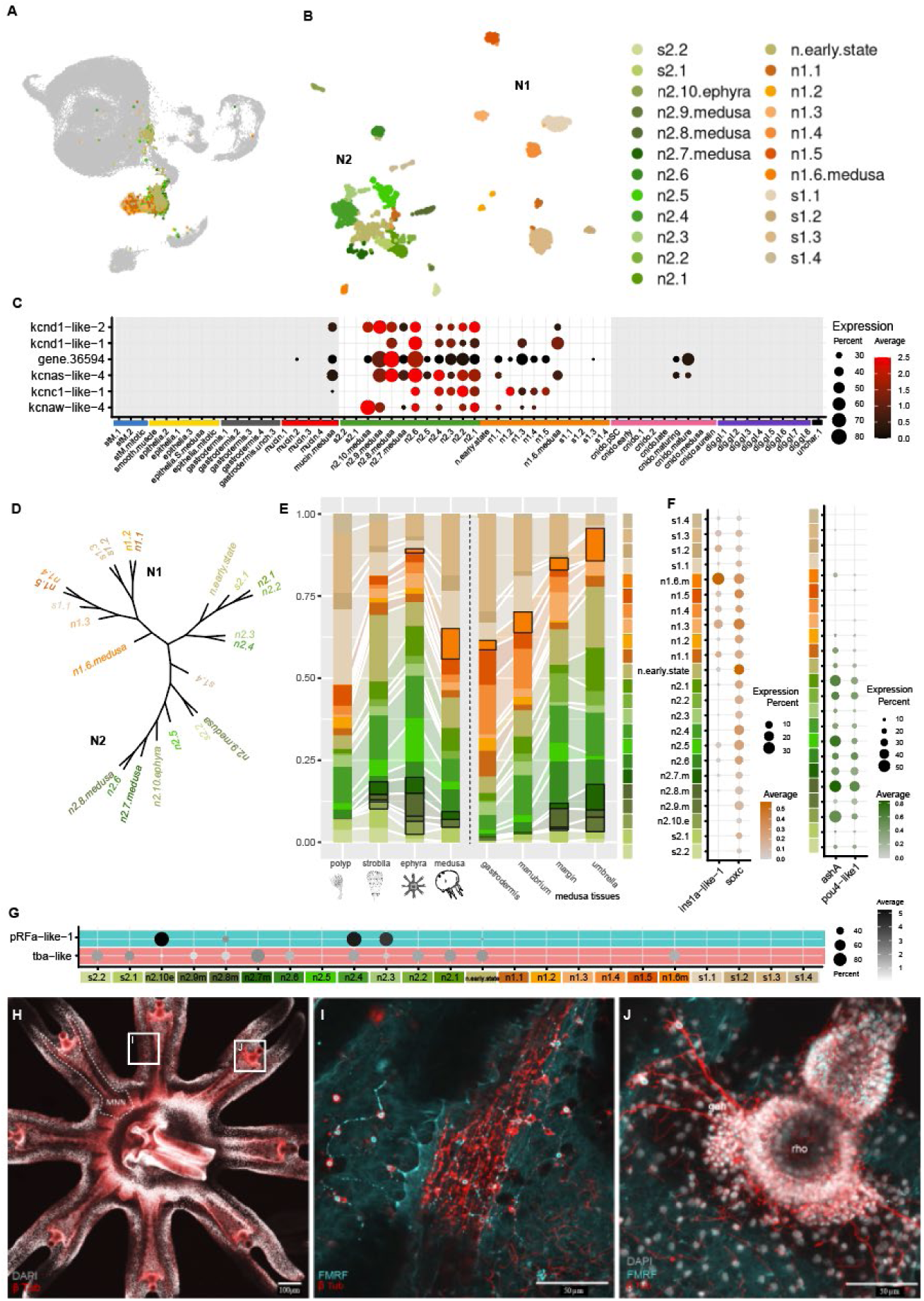
Neural cell complement includes two distinct classes, with medusa-specific subtypes in each. (**A**) Location of neuroglandular subset within the full dataset. (**B**) UMAP embedding of the neural subset (**C**) Expression of select voltage-dependent potassium channels across the full dataset. (**D**) Cell state tree of all neural sub-types (**E**) Distribution histogram across all samples, medusa-specific cell clusters that are absent from all polyp samples are highlighted with black outline. (**F**) Two classes can be recognized by differential expression of transcription factors. Class 1 cells express *insulinoma* (*ins1a*) (orange left) and class 2 express *achiote-scute* (*ashA*) and pou4 (*pou4-like1*) (green right). (**G**) Dotplot expression plots illustrating N2-specific ⍺-Tubulin (*TBA-like*) and precursorFMRFamide (*pRFa-like-1*) expression. (**H**) An anti-βTubulin antibody stains the motor nerve net of the ephyra. (**I**) Co-staining of anti β-Tubulin and anti FMRFamide visualises cells that are positive for both antibodies, possibly corresponding to cells expressing both tubulin and FMRFamide. The ephyra lappet displays condensed regions of the β-Tubulin positive motor nerve net aligning with the radial canal of the lappet. (**J**) Network of the rhopalia nervous system positive for β-Tubulin. DNN = diffuse nerve net, gan = ganglion, lap = lappet, man = manubrium, MNN = motor nerve net, rho = rhopalia.

In contrast, neurons of class 2 comprise most neuron types within the medusa umbrella and margin samples and include five neuron clusters that first appear between strobilation and release of the ephyra (**Fig. 4E**). Within the polyp, only the poly-RFamide class 2 neurons are abundant (“*n2.3*” and “*n2.4*”: **Fig. 4G**), and all other class 2 neural types are under-represented or absent (**Fig. 4E**). *Ins1*-negative class 2 neurons express both the bHLH transcription factor *ashA* and a *pou4/brn3* ortholog (**Fig. 4F**) that is shared also with the cnidocytes, similar to *Nematostella* (11,12). Class 2 neurons (**Fig. 4E**, shades of green) increase in abundance during the transition to the medusa form, including the generation of four additional medusa-specific class 2 neurons. Three are present already in the early strobila and are localized to the medusa margin and umbrellar samples (“*n2.7.m*”, “*n2.8.m*”, and “*n2.9.m*”), whereas “*n2.10.e*” appears only in the strobila and ephyra samples. The candidate specification marker for neuron “*n2.7.m*” is an atonal-like bHLH family transcription factor, *atoh8d*. We detect an expansion of the ATOH8 gene family, with detectable expression for 4 gene models. *atoh8b and atoh8c* are expressed in all class 2 neurons, while others show expression restricted to medusa-specific cell types: *atoh8d* (“*n2.7.m*” neuron) and *atoh8a* (“*mucin.medusa*”) (**Fig. S5, S8**).

We next sought to connect the single cell transcriptomic data to the anatomy of the nervous system. The scyphozoan nervous system is composed of three distinct parts: a diffuse nerve net (DNN) that covers the entire animal and is comprised of four anatomically distinct neural structures, the giant-fiber or motor nerve net (MNN) that is comprised of larger bipolar neurons that overlie the muscle fields, and the marginal ganglia or rhopalia that is located at the tips of the ephyra and later the margin of the medusa (49). We find that class 2 neurons all express elevated levels of specific alpha- and beta-tubulins (*tba1-like3* and *tbb-like-1*; **Fig. 4G**). Anti-β-tubulin antibody staining highlights concentrations of the MNN centrally located within the lappets of the ephyra ((23), **Fig. 4H**). Morphologically, this is the territory that includes the radial canals and overlying muscle fields of the ephyra. FMRFamide antibodies stain a subset of β-Tubulin positive portions of the medusa nerve net (**Fig. 4I**). Whereas the β-Tubulin antibody recognizes cells predominantly found near the striated muscle field and ganglion cells projecting out of the rhopalia, FMRFamide positive neurons are more scattered across the ectoderm (**Fig. 4J**). Some FMRFamide positive neurons are also found within the distal-most portion of the rhopalia (**Fig. 4J**). This distribution suggests that the β-Tubulin antibody targets predominantly the MNN, whereas the FMRFamide positive neurons form part of the DNN as previously reported (22). Specific tubulin-paralog expression within the class n2 neurons suggests that these two genes are translated into proteins recognised by this commercial β-Tubulin antibody.

### Muscle profiles correspond to smooth and striated fibre types

Striated muscle cells are known to be a specific feature of the medusa stage (24–26), yet a more detailed molecular description of muscles in jellyfish is lacking. To this end, we investigated the molecular muscle profiles and described the dynamic changes of muscle cell populations during metagenesis in *Aurelia coerulea*. One of the main partitions in our single cell transcriptome dataset defined by a set of muscle-related structural genes appears at the strobila stage (**Fig. 2C**). This cluster contains a large cellular contribution from the umbrella library, which contains the field of striated muscles. By contrast, this cluster is missing from the manubrium and gastrodermis tissue libraries, which only display smooth muscles (**Fig. 2C**). This suggests that this cell cluster represents the striated musculature. *Aurelia* muscles show a clear molecular signature specifically enriched in muscle related structural genes such as myosin, actin, and actin-binding proteins (**Data S.1.3**). We generated lists of muscle-type specific variable genes that are up-regulated in at least 10% of cells in either muscle cluster, and detectable in less than 40% of any non-muscle cluster. While smooth and striated muscles are clearly distinguishable by the differences in their molecular content, a substantial fraction (∼28%) of both muscle profiles consists of genes with no identifiable sequence identities that would indicate gene orthology (**Data S1.11**). We also identified a distinct transcription factor profile for striated muscle cells (**Data S1.3b, S1.4b**). One of the highest expressed transcription factors in striated muscle cells of *Aurelia* belongs to the bHLH family and shows sequence similarity to the anthozoan cPATH family (50). Additional transcription factors expressed in this cell type include two uncharacterized zinc fingers (klf1-like-1 and zn787-like-1), a homeodomain family transcription factor of uncertain affinity (gscb-like-1), and two *otx* paralogs, *otx1a*, [aka *otx1* (29)], and *otx2* (**Fig. S9; Data S1.3b, S1.4b**). *Otx1a* expression was previously shown to overlap with striated muscle fields, however it was inferred to define the overlying neuroectoderm of the motor nerve network (29). The current data does not exclude neural expression, as this gene is also detected within some neural populations (**Fig. S9**).

We assessed the anatomic location of the muscle fields by phalloidin staining in *Aurelia* polyps, strobilae and ephyrae (**Fig.5**). Polyps have three distinct smooth muscle fields (**Fig. 5A,B-G**): the radial muscles of the oral disc (**Fig. 5D**), the longitudinal tentacle muscles (**Fig. 5E**), and the longitudinal retractor muscles that run along the body column (**Fig. 5F,G** (35)). During strobilation, fragments of the polyp retractor muscles are retained in the early ephyra (**Fig. 5J** (35)). Striated muscles appear coronally around the oral disc, oriented radially along the lappets of early detached ephyra (**Fig. 5L-N**). At the tips of the lappets, the border of the coronal muscle, and at the base of the manubrium, fibres show a mixed organization of smooth and striated myofibrils (**Fig. 5O,P**). These findings corroborate previous studies that used light-(26) or electron microscopy (24,25).

**Fig. 5.**
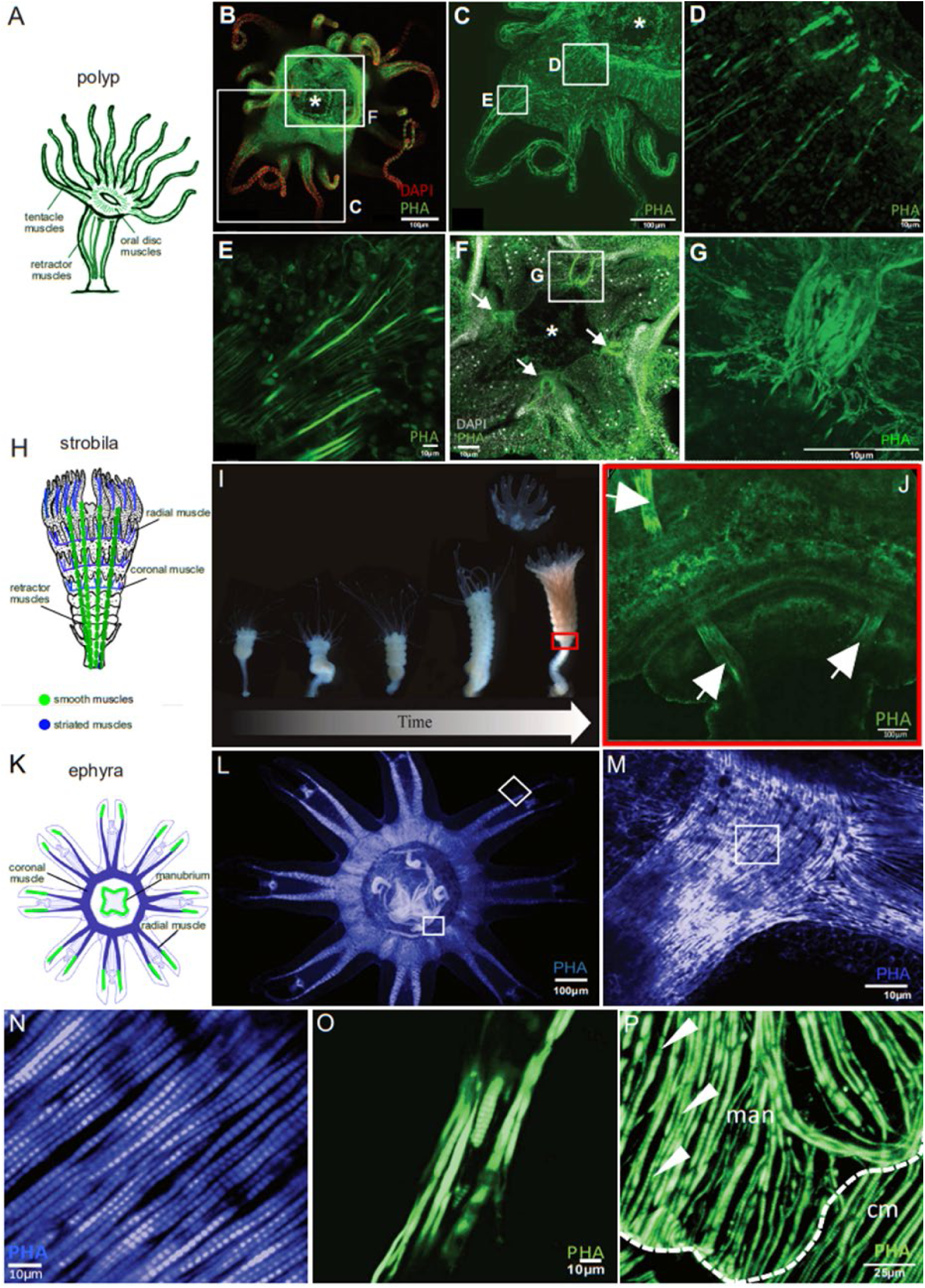
Muscle territories in *Aurelia coerulea*. (**A**) Schematic representation of the different smooth muscle territories in the polyp. (**B-G**) Phalloidin stains actin-rich muscle fields in the animals. Arrows: retractor muscles. (**H**) Schematic of a strobila illustrating the ephyra striated muscle fields (blue) and the fragmented polyp retractor muscles (green) (**I**) Consecutive stages of strobilation starting with a normal polyp to the first detaching ephyra. The presence of polyp retractor muscles in the undifferentiated foot of the strobila and the lowest ephyra in the stack. (**K**) Schematic representation of striated muscle fields in the ephyra. (**L-P)** Phalloidin stain visualizes coronal and radial striated muscles in the ephyra with some spots that show a mixture of smooth and striated myofibril organization. Arrowheads: myofibrils. man = manubrium, cm = coronal muscle.

We next compared expression patterns expected from our single cell data with the phalloidin-based anatomy of smooth and striated muscles. As expected, several genes were shared between the smooth and striated muscle cluster (**Fig.6E**), while others were highly specific to either smooth (**Fig.6C,D**) or striated muscle cluster (**Fig.6P**; **Data S1.11**). Different *calponin* paralogs show distinct expression in the different muscle types (**Fig. 7A**). For example, *calponin1* is specific to the smooth retractor muscle of the polyp and no other subpopulation of the smooth muscle type (**Fig. 6A-C**). At the strobila stage, expression of *calponin1* is still visible in fragmented retractor muscles, consistent with the single cell expression profile (**Fig. 6F**). By comparison, *mrlc2* expression marks the locations of all smooth muscle populations in polyps including tentacle muscles, radial muscles of oral disc and retractor muscles of the body column (**Fig. 6D,E**).

**Fig. 6:**
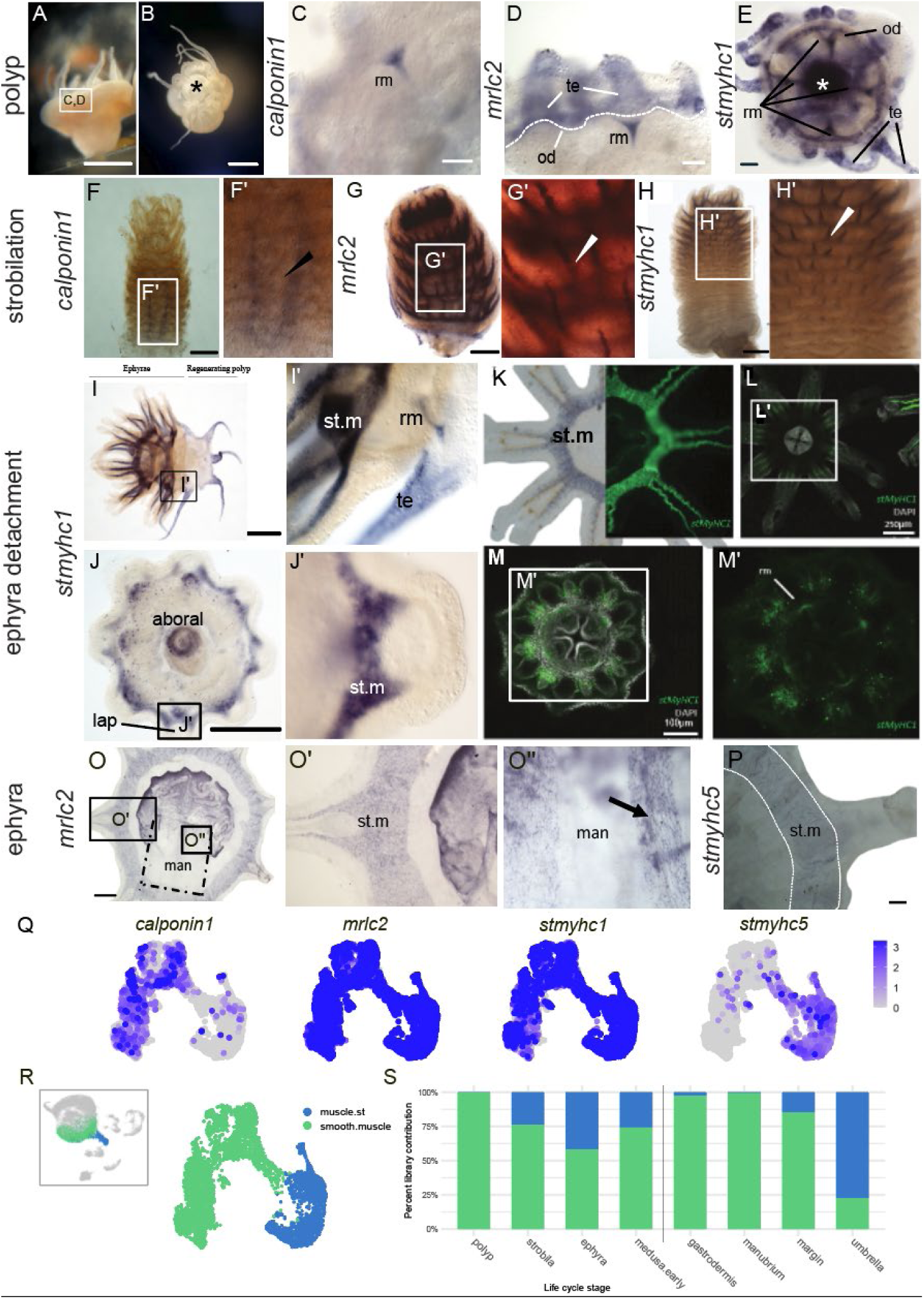
Gene expression profiles of smooth and striated muscle marker genes during the transition from polyp to ephyra. (**A,B**) Brightfield images show a lateral (A) and oral (B) view of an *Aurelia* polyp. Asterisk: mouth; scalebars: 1mm. (**C:E**) *in situ hybridization* in polyps. Expression in the different smooth muscle types of the polyp: *calponin1* (C)*, mrlc2* (D) and *stmyhc1* (E), scalebar: 200 μm. (**D:H**) Expression in strobila. *calponin1* (F) in the retractor muscle, *mrlc2* (G) and *stmyhc1* (H) in the developing striated muscle fields Arrowhead: retractor muscle; scalebar: 500μm. (**I:M**) *stmyhc1* expression. (**I,K**) in the advanced strobila. scalebar: 500μm (**K-M**) in the detached ephyra. (**O**) *mrlc2* is detected in coronal muscles and in the manubrium in the ephyra. Scalebar: 500μm. (**P**) *stmyhc5* labels the coronal muscle in the ephyra. Scalebar: 500μm. (**Q**) feature plots of all marker genes on the muscle specific subset (**R**) reference UMAP of whole dataset (left) subset (right) (**S**) Distribution plot of muscle types across the different *Aurelia* life stages (left) and medusa tissues (right). st.m = striated muscles, rm = retractor muscles, man = manubrium, te = tentacles, od = oral disc, lap = lappet.

**Fig. 7:**
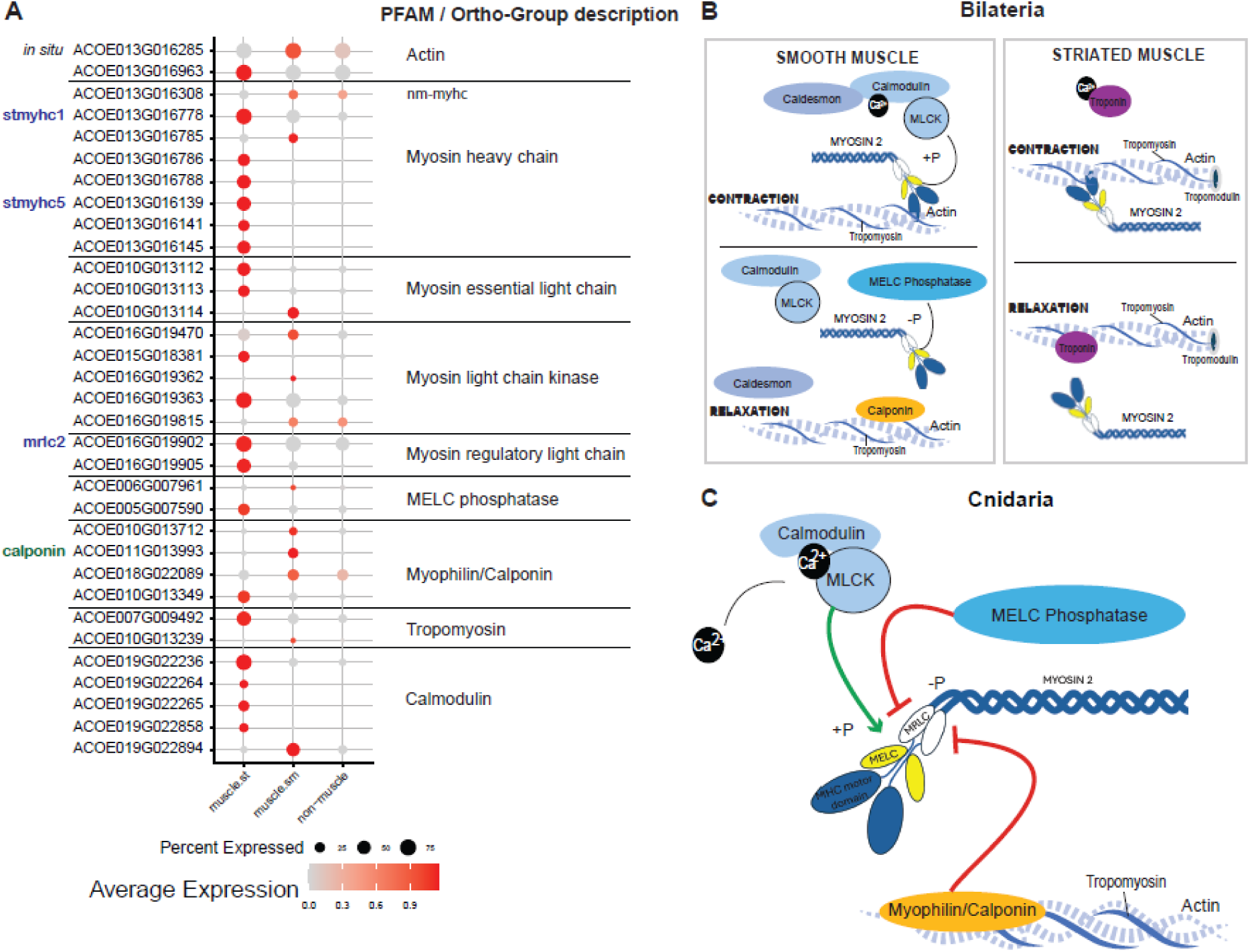
Shared contractile complex between smooth and striated muscle cells. (**A**) Dotplot of gene expression for muscle-type specific gene sets. (**B**) Schematic shows main molecular components for myosin-based contraction in bilaterians. (**C**) Hypothesis of muscle contractile apparatus in Cnidaria.

### The emergence of striated muscles during metagenesis from polyp to medusa

Strobilation marks the transition from the polyp to the medusa and hence the emergence of striated muscles in the developing medusa. Myosin heavy chain genes duplicated in a unicellular ancestor, giving rise to the smooth/non-muscle myosin heavy chain (*myhc*) and a paralog, called striated type myosin heavy chain (*stmyhc*) because it is specifically expressed in bilaterian striated muscle (51,52). Notably, in the sea anemone *Nematostella vectensis, stmyhc* is strongly upregulated in the fast-contracting smooth muscle (50,51,53), suggesting that this variant does not convey striation but rather is associated with fast contraction. In *Aurelia*, we detect seven *stmyhc* paralogs expressed within the muscle clusters (**Fig. 7A**). We examined the expression profile of two of these models (*stmyhc1* and *stmyhc5*) which are expressed in striated muscle cells. While *stmyhc5* is restricted to striated muscles, *stmyhc1* is also expressed in the smooth body column retractor muscles and tentacle muscles of the polyp (**Fig. 6E,H**). In the strobila, *stmyhc1* expression remains expressed in both intact (**Fig. 6I**) and fragmented smooth polyp retractor muscles (**Fig. 6L,M**), consistent with previous studies showing that polyp retractor muscles persist through strobilation until disappearing in early detached ephyra (26). Notably, in the regenerating polyp *stmyhc1* is again expressed in the smooth muscles of the newly forming polyp tentacles, reflecting its role in tentacle smooth muscles (**Fig. 6I**, (50)). In the strobila and ephyra *stmyhc1* expression is also detected in developing coronal- and radially-oriented striated muscles (**Fig. 6I,J,K**). In contrast, the *stmyhc5* paralog is restricted to the mature striated muscles in the late ephyra (also called “metaephyra”), an intermediate stage between ephyra and juvenile medusa (54), (**Fig. 6P-S; Data S1.10**), suggesting that it plays an important role in the fast-contracting striated muscles of the jellyfish. Like *stmhc1*, *mrlc2* (*myosin regulatory light chain 2*) is also shared between the two muscle subtypes, raising the interesting possibility that striated muscles arise from a smooth muscle progenitor state.

### Principles of muscle contraction and sub functionalization of paralogs

Both *Aurelia* muscle subtypes express a set of genes that comprise a core ancestral contractile complex that includes *calmodulin*, *myosin light chain kinase (mylk)* and *tropomyosin* (*tpm*) together with *myosin heavy chain*, *essential*, and *regulatory light chains* (*myhc, mel, mrlc*) (**Fig. 7A**; **Data S1.11**). Vertebrate striated muscle differs from smooth muscle in that it uses *troponin* to confer calcium-sensitivity to striated muscle actomyosin ATPase activity. As cnidarians do not have bona fide Troponin homologs, the regulation of contraction relies on myosin light chain kinase (MYLK) and the phosphorylation of myosin light chain (MELC) in both smooth and striated muscle. This regulatory gene complement is shared with bilaterian smooth muscles (**Fig. 7B,C**), suggesting that this represents the ancestral mode of contraction regulation (51,55).

Notably, almost all the genes coding for these proteins have undergone multiple gene duplications as evidenced by the multiple paralogs present in the genome. These paralogs are often clustered as tandem repeats, as indicated by sequential gene model numbers (**Fig. 7A**). In addition to a set of shared genes expressed in all muscle subtypes (**Data S1.11**), smooth and striated muscles additionally express distinct sets of paralogous core muscle proteins across the two muscle types (**Fig. 7A**), suggesting a process of subfunctionalization following gene duplication. For instance, we find four myosin essential light chain paralogs, of which two are differentially expressed in smooth and two in striated muscles (Fig. 7A). In almost all cases the expression of the paralogs is divided between the two muscle types. Differences in the molecular features of these paralogs may explain the differences in contraction speed and force. This is remarkably similar to the subfunctionalization of paralogs of muscle proteins in *Nematostella* (50).

## Discussion

### Single cell transcriptomic atlas reveals distinct cellular subtypes associated with the formation of medusa in the moon jelly

Here we present a single cell transcriptome characterisation of the metagenesis in a scyphozoan, the moon jelly *Aurelia coerulea,* in which we document an overall increase in gene detection of approximately 10% during formation of the medusa. Other studies have examined differential gene use from bulk RNA extracts from medusa versus polyp stages and clearly demonstrate changes in the overall gene representation across the life cycle (20) but have found no evidence for taxon-specific genes in either stage (18), or even in the composition of reconstructed proteome or transcriptomes from polyps compared to medusa (56). While we find only one polyp-specific digestive cell type, we document several distinct cell types associated with the generation of the medusa stage that become detectable in the strobila but are absent from all the polyp samples. Apart from the striated muscle, which is an abundant cell type of the medusa, these medusa-specific cell types comprise a small fraction of the overall tissue composition and share transcriptomic similarity with related cell types that are present across the life cycle.

Of the cell clusters that are absent from the polyp stage, five of these are neurons. We can tentatively associate these neural subtypes with components of the nervous system known to arise in association with the formation of medusa-specific tissues, including the rhopalia, motor nerve net, and striated muscle (8,22,57). We note extensive parallels in the complement of neural subtypes with the organisation described in the sea anemone, particularly the partitioning into two major neural classes. This division is similar to anthozoans, where Steger *et al*. (11) identified mutually exclusive *insulinoma* positive and *POU4* positive neural trajectories in *Nematostella*. In *Nematostella* the *ins*-positive neuron population is more abundant and possesses a greater number of subtypes, including the sensory/secretory “S”-class of largely uncharacterized cell types. It is this n1 subset that also predominates the polyp stage of the lifecycle in *Aurelia*. This contrasts with the distribution of n1 and n2 class neurons in the freshwater hydozoan polyp *Hydra vulgaris*, of which only three of the fifteen sub-types are of the *ins*-positive n1 type (“ec2”, “en2”, and “en3”: **Fig. S8D;** (58)). Similarly in the *Clytia* medusa only a fraction of one of the three identified neuron groups (neuron cells “A” (16) have *INSM* reads and thus could be considered type 1 neurons as defined here. The digestive gland cell populations in both *Aurelia* and *Nematostella* express orthologs of the achaete-scute family (*ashC*) (11), suggesting an ancestral role in specifying secretory cell types for this gene. *AshC* is an ortholog of the *ascl345* family in vertebrates, which is expressed in the skin (*ASCL4*), salivary glands (*ASCL5*), and teeth (*ASCL6*) [UNIPROT database (59)]. Orthologs of *ashB* and *ashD* are expressed in the neurons in both *Nematostella* (11,12) and *Aurelia,* and both are homologs of the Achaete-scute complex, notable regulators of neurogenesis in bilaterians (60). AshA does not have a mouse or human ortholog, but is present within nematodes, where it is a key regulator of a nociceptive neuron identity (61). In *Aurelia,* the *ashA* ortholog is restricted to the class 2 neurons, while in *Nematostella ashA* expression is found in both types (11,12,62) and *ashD* is restricted to INS-positive (n1) neurons in both species. The *ins*-negative populations in both species express *pou4* orthologs, also called *brn3* (29), that is also expressed within the cnidocyte lineages and thus further supports claims of a close relationship between cnidocytes and *insulinoma*-negative/*pou4*-positive n2 neurons (11,12,48). This strongly suggests that the radiation of achaete-scute genes not only preceded the split of cnidarians and bilaterians, but also that the paralogs were implemented in the diversification of the neuro-glandular lineages across the Cnidaria. Further, while we can distinguish two neuron molecular classes of neurons (**Fig.4 A-E**), our results do not support a 1:1 relationship between these two neuron classes and the morphological and functional distinction of the previously identified medusa DNN and MNN (22,23,28,29). While tyrosinated-tubulin and beta-tubulin antibodies have been shown to label the MNN ((22) and this study), neural-specific tubulins are expressed in neurons of the n2 class (**Fig. 4F**). However, the neuropeptide *fmrfamide* is expressed in the DNN (22), is also restricted to neurons of the n2 class (**Fig.4 F-D**). Therefore, it seems that the DNN and MNN both contain class n2 neurons. The distribution of class n1 neurons is currently unknown.

The other major neuroglandular cell type family is the cnidocytes. Our single cell transcriptome data suggests the presence of two major capsule types in the tissues of *Aurelia coerulea* (**Fig. 3**). While the description of the *Aurelia aurita* cnidome identified six different types of cnidocytes, only two capsule types, a-isorhizas and euryteles strongly dominate the tissue of this species (37). Given the close phylogenetic relationship, this is likely also the case in *Aurelia coerulea* and the abundance of these two capsule types might be reflected in the resulting scRNA sequencing data. Strikingly, the comparison of the developmental trajectory of cnidocyte development between *Nematostella vectensis* and *Aurelia coerulea* reveals a stunning level of conservation of gene regulation, despite significant morphological differences in cnidocyst shapes (11,63,64), suggesting that while cnidocyte diversification has occurred in the different lineages, the genetic regulation of cnidocyte development predates the split of medusozoans and anthozoans.

### Transcription factor family expansions may be related to the generation of novel medusa cell types

Recently, we demonstrated a sea anemone specific expansion of an orphan bHLH family that correlated with the diversification of muscle cell types (50). Expansions of other transcription factor families in a similar manner could be related to generation of novel cell types (63). In samples from strobila, ephyra and medusa we found four different medusa restricted neuron subtypes (“*n2.7.m*”, “*n2.8.m*”, “*n2.9.m*” and “*n2.10.e*”). These four neuron types likely correspond to sensory cells of the rhopalium and putative motor neurons that innervate the striated swimming musculature, as these tissues are medusa-specific and first develop within the strobila. Published expression data further supports this assumption: *Pit1* expression has been reported in putative sensory neurons of the developing rhopalia (29) and is expressed in “*n2.5*”. Neuron “*n2.10.e*” expresses the homeodomain transcription factor *otx1c* as well as poly-RFamides (pRFa) (**Fig. 4F**). We also identified a set of *insulinoma*-negative/*pou4/brn3*-positive n2 neurons, which likely represent the *pou4/brn3* positive population identified by Nakanishi *et al.* (29). The medusa-specific cell types identified here also demonstrate cell type specific expression of expanded transcription factor families. A good example is the expansion of the ATOH8 gene family. While the gene models *atoh8b and atoh8c* are expressed in neurons of both sessile and free-swimming life-stage, *atoh8d* is expressed exclusively in medusa specific neuron “*n2.7.m*”. The ortholog of *atoh8d* in *Nematostella* is expressed in the two gastrodermal N2 class neurons (“*N2.g1*” and “*N2.5*”; (12)), while in *Hydra* it is expressed in n2-class ectodermal neuron 3 subtypes (“*ec3A”,* “*ec3B”,* “*ec3C”*; **Fig. S8**). *Atoh8c* and *atoh8d* appear to be cnidarian-specific paralogs (**Fig S8E**). This suggests the “*mu.medusa*” cell type may be a novel cell type that arose after gene duplication, as has been proposed for anthozoan muscle (50), whereas “*n2.7.m*” may be a cell type ortholog of the n2 gastrodermal neurons in the anthozoans. This is an intriguing observation as the gastrodermal neurons lie near the longitudinal parietal muscle of the inner cell layer (65). We cannot at this time distinguish between gastrodermal and ectodermal neurons in the *Aurelia* dataset. The *atoh8d* ortholog is expressed in a type n2 ectodermal neuron (Ec3N) in the freshwater *Hydra* polyp, where inner gastrodermal and outer ectodermal neurons have been distinguished (15,58,66), **Fig. S8D**). Other examples are *myc* genes that also appear to be expanded within the cnidarians, at least in *Nematostella* (6 copies (67)) and scyphozoans (7 or 8 copies; **Fig. S6A**). While four paralogs have been reported from the hydrozoan *Hydra* (68), only 2 have been investigated (69–71). Similarly, only 2 paralogs are reported from another hydrozoan (72). Interestingly, of the two *hydra myc* genes, one is expressed in the cnidocyte lineage, while the other is expressed in the interstitial stem cell lineage ((15), **Fig. S6B**). The *myc* expansion in *Nematostella* and *Aurelia* are used in parallel ways during cnidocyte development, and orthologous pairs of *Nematostella* and *Aurelia* myc genes are similarly expressed in the early and mature phases of cnidocyte differentiation (**Fig. 3E:F, Fig. S6**). These data support the hypothesis connecting gene duplication with cell type expansion (50,51,73–77), suggesting that at least some of the medusa-specific cell types identified here may represent synapomorphies of the Medusozoa, or perhaps even the Scyphozoa.

### Shared core contractile machinery between smooth and striated muscles indicates a gradual maturation of smooth fibres into the striated phenotype

Striated muscles in the subumbrella enabled jellyfish to evolve a unique swimming behaviour and opened a new ecological niche. To infer the evolution of this muscle cell type, we investigated muscle cell transcriptomes across different stages of the *Aurelia* life cycle, including both striated and smooth types. The striated muscle population is the largest medusa-specific cell population with a distinct transcriptomic profile. The smooth muscle transcriptomic profile shares features with the epidermis and is recovered as a cell state within this larger population, probably reflecting its myoepithelial nature **(Fig. 2A)**. While a small set of genes are shared across the two muscle phenotypes **(Fig. 7A, Data S1.11)**, we also report the expression of paralogues specific to either smooth muscles **(Fig. 6C,Q**, **Fig. 7A)** or striated muscles **(Fig. 6P,Q**, **Fig. 7A)**. The shared expression of muscle-related structural genes is most likely part of a core contractile machinery that is present in both smooth and striated muscles (**Fig. 7B,C; Data S1.11**). Tanaka *et al.* (55) linked some genes expressed in *Aurelia* striated muscles to a bilaterian smooth muscle contractile apparatus. This core contractile complex is regulated via phosphorylation of *myosin regulatory light chain* (*mrlc*) by *myosin light chain kinase* (*mylk*) in the presence of Ca2+, which is known to regulate bilaterian smooth and non-muscle cell contraction (**Fig. 7B**) (55,78,79). Main players in this smooth contraction mechanism are *calmodulin*, *mrlc*, *melc* and *mylk*, which are all expressed in both *Aurelia* muscle types identified from our single-cell data (**Fig. 7A; Data S1.11**). We found *striated myosin heavy chain* paralogs expressed in all muscle populations whereas other paralogs are specific to the striated muscle state (**Fig. 6P, 7A**). This suggests a role for the various *myhc* paralogs in differentiating muscle architecture. In vertebrates, switches of different isoforms of *stmyhc* genes during maturation of striated muscle cells between embryonic and perinatal stages have also been reported (80). In addition, we found other structural protein gene families resulting from one or several gene duplications, where the paralogs became specifically expressed in either smooth or striated muscle. We suggest that the subfunctionalization of paralogs of structural protein genes may be the cause for the physiological differences of the two muscles with respect to contraction speed and force. A similar subdivision of paralogs was found in the sea anemone *Nematostella vectensis*, where two fast and two slow contracting smooth muscle cell types exist (50).

At this point, it is still unclear whether striated muscle cells derive from a distinct progenitor cell population or from a subset of smooth muscle cells. We observe individual smooth fibres located at the base of the developing manubrium that show striation at some locations along the fibre (**Fig. 5O,P**), raising the possibility that smooth and striated fibres can be found in the same cell. This phenomenon is known from the vertebrate heart muscle, where non-striated pre-myofibrils gradually undergo sarcomere assembly during myofibrillogenesis (81). In scyphozoan species like *Aurelia coerulea* this could be the case during metamorphosis from the polyp to the medusa form. However, future work is required to further provide experimental evidence for this scenario.

## Material and Methods

### *Aurelia coerulea* (formerly sp.1) culture

The lab strain was obtained from the collection of Dr. Harms (University of Hamburg) approx. 30 years ago. Genomes of several *Aurelia* strains have recently been published. Transcriptome reads were first mapped to all available *Aurelia* genomes (18,82), with gene annotations extended by 1000bp towards the 3’ end, or until reaching the next gene model in the same orientation (83). We were able to align 90% of the reads from our libraries to the genome of the San Diego aquarium strain (18), compared to only 40% aligning to the atlantic strain or 84% to the pacific/Roscoff strain (82). CO1 gene trees cluster the Roscoff and SanDiego aquarium strain together with *Aurelia sp.1*, which has now been identified as *Aurelia coerulea* (84). From these data we identify our laboratory strain as a member of the *coerulea* species. *Aurelia* scyphistomae were grown on small petri dishes placed in 80ml glass containers filled with ASW (artificial sea water) with the salinity adjusted to 35‰. Polyp cultures were fed with *Artemia* nauplii once a week. Containers were cleaned from algae approximately once a month. Strobilation of polyps was induced through either a shift in temperature of about 2-5°C or an 24h treatment with 10µM Indomethacin (38). Freshly liberated ephyrae were transferred to a 5L beaker. Water motion was induced via plastic propellers attached to a small 12V engine. Ephyrae were fed once a day with freshly hatched artemia nauplii. Small jellyfish were transferred to a 30L Kreisel Tank and daily fed with *Artemia* nauplii, which were incubated in Selco S.presso (INVE) and diluted isochrysis extract overnight to enrich live food with nutrients. Kreisel Tanks were cleaned once a week.

### Single-cell dissociations

Magnesium-calcium-free artificial seawater (MCF-ASW) was prepared according to (85). Scyphistomae, strobilae, ephyrae, or juvenile medusae were collected separately into 1.5ml tubes, washed 2x with filter-sterilized artificial sea water (FSASW) and incubated in MCF-ASW, followed by incubation in 300μl Collagenase/Dispase Blend I (Merck) dissolved in 35‰ FSASW at room temperature (RT). Mechanical force via pipetting was used to disrupt the rest of the tissues and obtain single-cell suspensions after 30 minutes of enzymatic digestion. The suspension was monitored for cell clumps using a compound microscope. To stop the digestion once cell clumps were fully dissociated, the suspensions were put on ice and 0.1% bovine serum albumin (BSA) was added. Cell suspensions were transferred to a pre-coated (0.2% phosphate buffered saline with 0.1% tween (PTW) followed by extensive washing) 1.5ml tube and pelleted for 5 minutes and 0.4rcf using a pre-cooled centrifuge (10°C). The supernatant was removed, and cells were resuspended in 100μl of BSA (0.1% final concentration) and 100ml MCF-ASW. After resuspension samples were filtered through a Flowmi-filter (40μm). Cell recovery and viability were measured with a Nexelcom cell counter upon incubation with 10μM Fluorescein and Propidium iodide. Finally, quality single-cell suspensions were diluted to 1000 cells / uL and loaded according to kit protocols into microfluidic chips with the target cell capture of 7000 cells (10X Genomics, three-prime RNA, vs. 3). Tagged single cell RNA was then processed into a sequencing library following the manufacturer protocol. The resultant single-cell libraries were sequenced with the Illumina platform according to the 10X genomics specifications for paired end sequencing. Sequencing statistics for all libraries are included in supplementary **Data S2.1**

### Transcriptome

Chromosome level genome assembly was downloaded from NCBI (GCA_039566865.1). Genomic scaffolds (22 chromosomes and shorter unplaced fragments) were sorted according to their lengths and renamed to scaffold_1 to scaffold_22. Species-specific gene prediction was done using AUGUSTUS v3.5.0 (86) was trained by using transcriptomic data from planula, polyp, strobila and jellyfish stages of the laboratory strain "Roscoff" (*A. coerulea*) and the animals from Kuyukushima aquarium (see NCBI TSA accessions GHAI00000000 and GHAS00000000). These transcriptomic data were mapped against the genome assembly and the resulting hits were converted into hints for exons and exon-intron boundaries used by AUGUSTUS gene prediction program. Additionally, an inhouse transcriptome assembly was aligned to the annotated genome using minimap2 (-x splice, -L, -c –MD, –eqx). The paf alignment was then filtered to retain high quality primary alignments (MAPQ >= 30) that were not present in the initial annotation. The PAF alignments were converted to GTF files using an inhouse script (see code availability). Gene model nomenclature adopted here follow the scheme ACOE(species)0##(chromosome)G######(gene number) wherein the gene number follows the order along the chromosome. Final transcripts were functionally characterised using different databases including Pfam, Uniprot, NCBI NR, InterproSites, TMHMM, SignalP and eggnogmapper. Transcription factors were identified and classified using the Animal Transcription Factor Database version 4. (https://guolab.wchscu.cn/AnimalTFDB4/ (87–90)). Putative names were assigned based on sequence homology to Uniprot best blast hit or NCBI NR best blast hit. A look up table to convert between the gene models used here, the nomenclature provided, and the previously published gene models is available in supplementary material **Data S2.2.**

### Phylogenetic Trees

Putative bHLH domains were identified using HMMER (3.4) with default parameters from the following proteomes: *Amphimedon queenslandica (NCBI: GCF_000090795.2), Hydra vulgaris (*https://research.nhgri.nih.gov/HydraAEP/download/index.cgi?dl=pm*), Aurelia coerulea* (this study: available from https://genome.ucsc.edu/h/GCA_039566865.1)*, Magallana gigas (UNIPROT: UP000005408), Branchiostoma floridae (UNIPROT: UP000001554), Mnemiopsis leidyi (*https://research.nhgri.nih.gov/mnemiopsis/download/download.cgi?dl=proteome*), Caenorhabditis elegans (UNIPROT: UP000001940), Nematostella vectensis (*https://simrbase.stowers.org/starletseaanemone*), Cassiopea xamachana (*https://genome.jgi.doe.gov/portal/pages/dynamicOrganismDownload.jsf?organism=Casxa1*), Owenia fusiformis (UNIPROT: UP000749559), Drosophila melanogaster (UNIPROT: UP000000803), Rhopilema esculentum (*https://www.ncbi.nlm.nih.gov/datasets/genome/GCF_013076305.1/*), Ephydatia muelleri (*https://spaces.facsci.ualberta.ca/ephybase/*), Sanderia malayensis (NCBI: GCA_013076295.1), Stylophora pistillata (UNIPROT: UP000225706), Exaiptasia diaphana (UNIPROT: UP000887567), Homo sapiens (UNIPROT: UP000005640), Tripedalia cystophora(*https://www.ebi.ac.uk/ena/browser/view/GHAQ01000000*), Hydractinia symbiolongicarpus (NCBI: GCF_029227915.1)*. Intra-species redundancy was reduced with CD-HIT (4.8.1) with a k-mer size of 5 and sequence identity threshold set at 97%. Multiple sequence alignment was performed with MAFFT (7.526) using the –localpair alignment option. Sequence alignment is found in supplementary Data **S3.** The phylogenetic tree was calculated with IQ-TREE (3.0.1) with automated model selection (-m MFP). Gene clades corresponding to the atoh8 (**Fig. S8**) and myc (**Fig. S6**) families were extracted from the tree.

### Single-cell analysis

Raw sequence reads were first processed through the cell ranger pipeline with forced recovery of 7000 cells and mapped using a custom mapping tool. Genes are named here according to their best blast hit whenever possible. UMI-collapsed count matrices were imported into R for further processing with the Seurat package (vs. 4.3) (91–93). Expression matrices from individual samples were first filtered to include only samples with at least 750 unique reads. Samples were further filtered for minimal genes (sample range: 350-500), and putative doublets were removed by eliminating outliers with high count values (sample range: 4000-40000). The data was then log normalised with a scale factor of 5000 reads (Seurat::NormalizeData), and top 2000 variable genes were identified for each sample independently (Seurat::FindMarkers). To avoid clustering according to cell cycle state, the resultant variable genes were filtered to exclude those involved in regulation of the cell cycle. Scaled gene expression of this combined set was used as input to the RunPCA function prior to data integration. Samples were then integrated with a reciprocal PCR approach, using the Seurat::FindIntegrationAnchors and Seurat::IntegrateData functions.

The resultant integrated dataset was then used for analyses of cell-state composition following a standard Seurat workflow. Briefly, the integrated data was scaled and used to identify the first 22 principal components with standard deviation of >2. These 22 components were used to generate a nearest neighbour graph (Seurat::FindNeighbors with annoy.metric = ‘cosine’, k.param = 10) from which cells were clustered with a resolution of 0.1. To visualise the clusters, all cells were projected in a two-dimensional space (UMAP). Each population was then re-analyzed separately, and distinct cell states and/or cell types from each population were annotated. Cell cluster identity was assigned semi-automatically as described in (11). Gene sets used to annotate the single libraries are found in Supplemental **Data S2.3:10**. These annotations were then imported into the full dataset for further exploration. To explore data from the perspective of the life history stage, samples of a similar stage were grouped together (polyp = 3, strobila = 2, ephyra = 2, medusa = 5). The cell state tree was calculated by first extracting the embeddings corresponding to the first 50 components computed with Seurat::RunPCA, generating a distance matrix with the euclidean method (stats::dist), and subsequently generating a neighbour joining tree with the ape R package (vs5.8: ape::nj). Upregulated marker genes for each cluster (or stage of the life cycle) were calculated with the Seurat ‘FindAllMarkers’ function, returning markers that show a fold-change equal or greater than 0.6, detected in at least 30% of the cluster, and having a p-value threshold of >=0.0001. Scripts for running all analyses and generating the figures found in the paper can be found on our GitHub page: Linketal.Ac.generate.data_REVISED.R; Linketal.Ac.analyse.alldata_REVISED.R; Linketal.Ac.analyse.subsets_REVISED.R; Linketal.Ac.generate.figures_REVISED.R : https://github.com/technau/AureliaAtlas.

### Comparison with *Nematostella*

For direct comparisons with *Nematostella*, we first ran OMA (45) with default parameters on the two transcriptome references. In total 4311 1:1 gene orthologs between the two species were identified (**Data S2.11**). A similar comparison using OrthoFinder (94) between Hydra and Clytia, both members of the Hydrozoa clade, found 5979 1:1 orthologs (66). OMA was preferred in this study over other available orthology databases because it outputs a high-confidence predicted 1:1 gene orthology list that can be used directly to combine multi-species data. The data matrices for the *Nematostella* single cell atlas of cnidocytes and neuroglandular cells is available at https://sea-anemone-atlas.cells.ucsc.edu under the “Nv2 Atlas/Cnidocyte subset” and “∼/All Neuroglands subset” subdirectories. To generate neural expression profiles for *atoh8* orthologs we first removed all “S”, “GD”, and “early” cell clusters, leaving only the set of putative neural profiles. To compare cnidocyte specification pathways, we removed the following cell clusters from the *Nematostella* dataset: “early”, “gast.1”, “planula.spir”, “planula.nem” and “planula.mat”. For the *Aurelia* cnidocytes we removed the clusters “early”, “medusa.1”, and “medusa.2”. The subset of each data matrix for the set of putative 1:1 orthologs (OMA) was extracted from both species and aligned using the harmony package (vs.1.2) (46). The cell state tree including both species was then calculated using 20 harmonized principal components and the neighbour-joining algorithm as described above. Support for the recovered nodes was evaluated with the ape::boot.phylo function with 1000 replicates. Script for preparing the data can be found on our GitHub page: Linketal.AcNv.cnidocytes.R : https://github.com/technau/AureliaAtlas

### Immunohistochemisty and phalloidin

Animals from different life-stages were relaxed at 4°C followed by a fixation step via 4%PFA at RT for 2h and washed two times in 0.2% PTW after fixation with a minimum washing time of 1/2h. Further, they were blocked using 3% (w/v) BSA (bovine serum albumin) for 2 minutes and incubated in a 1:50 PHA Alexa568/0.2% PTW and 1:1000 DAPI solution for 2h at RT in the dark. After staining they were washed 3x in 0.2% PTW, incubated in vectashield (mounting medium) at RT for 1/2h, mounted on glass slides and imaged using a confocal microscope (Leica SP8). In case of immunostaining, both the sample and the primary antibodies (βTubulin (Sigma Sc-5274) or FMRFamide (Millipore AB15348)) were blocked (1:300) in 0.1% FCS (fetal calf serum) for 1h at RT (sample) or at 4°C (AB). Afterwards the samples were incubated in the pre-blocked antibodies for 24h at 4°C and washed 2x with 0.2% PTW (1/2h/wash). Secondary antibodies (Goat α rabbit, life technologies; Goat α mouse/rat, Invitrogen; Alexa488) were blocked for 1h in FCS (1:1000) at 4°C and washed. The sample was incubated in the secondary antibodies for 20h (4°C). After incubation treatment followed the same steps as after PHA and DAPI stains.

### Cloning of genes and preparation of *in situ* hybridization probes

RNA was extracted from single-cells, starting with a digestion of animal tissues form different life-stages in 500ml of Animal Free Collagenase/Dispase Blend I (Merck) dissolved in 35‰ FSASW (filter sterile artificial sea water) prepared with DEPC treated water. Single cells were pelleted with 0.4rcf at 4°C for 5 minutes after digestion. The pellet was resuspended, and RNA extracted from single cells by following a standard Trizol extraction protocol. Primer sequences were designed using the primer3 website to amplify target genes of interest with the primer’s listed in the supplementary data file **Data S2.12**. Marker gene sequences amplified from cDNA obtained from extracted mRNA. Genes were cloned using a pJET1.2 blunt vector cloning kit according to the manufacturer. Plasmids were transformed into *E. coli* and bacteria grown on ampicillin treated agar plates afterwards. Minipreps were performed using a peqGold miniprep kit (VWR). WMISH probes were generated via a SP6 or T7 transcription kit (NEB) and prospectively labelled with either Fluorescein or DIG-UTP. WMISH or FISH were performed using a standard protocol with Hybe buffers based on Urea adapted to the needs of *Aurelia* tissues (95). Commercial fluorescein/POD and DIG/POD antibodies and buffers were used according to the manufacturer’s protocol (perkin elmer) for the performance of FISH.

## Supporting information

Supplementary Figures HTML

Differential Gene expression lists

Technical Information

Multiple sequence alignment

## Acknowledgments

The authors acknowledge the LISC (life science computer cluster) for providing infrastructure necessary for the bioinformatics, and CIUS (cell imaging and ultrastructure research) for technical support, equipment, and expertise. We thank the staff of the Tiergarten Schönbrunn (Vienna Zoo: Anton Weissenbacher, Roland Halbauer) and Haus des Meeres (Vienna Aquarium: Daniel Abed-Navandi) for sharing their expertise in cnidarian culture, and the UCSC Cell Browser (Maximilian Haeussler and Brittney Wick) for formatting and hosting these data for interactive exploration.

## Funding

This research was funded in whole or in part by the Austrian Science Fund (FWF) P34970 (UT) and P31018 (AGC). BW was funded by NIH/NIMH RF1MH132662, CIRM DISC0-14514 and NIH/NHGRI U24HG002371. For open access purposes, the author has applied a CC BY public copyright license to any author-accepted manuscript version arising from this submission.

## Author contributions

Conceptualization: OL, AGC, UT

Methodology: OL, JDM, BZ, AGC

Investigation: OL, SMJ, BZ, KJ, JK, JDM, AGC

Supervision: AGC, UT

Writing—original draft: OL, AGC

Writing—review & editing: AGC, UT, OL

## Competing interests

Authors declare that they have no competing interests.

## Data and materials availability

Raw sequence data will be deposited in the GEO database (https://www.ncbi.nlm.nih.gov/geo/), and the analysed data matrix is available on the UCSC Cell Browser (https://jellyfish-atlas.cells.ucsc.edu/). Scripts for analysing the data and generating the figures in this manuscript can be found on our GitHub page (https://github.com/technau/AureliaAtlas).

## Supplementary Material

### Figures

**Figure S1:** A) Total number of genes (nFeatures_RNA) and B) molecules (nCounts_RNA) recovered from all cells captured across life cycle stages. C) Distribution of medusa-specific genes across all cell types. The set of all medusa-specific genes were considered. For every cluster the gene is counted as expressed if reads are detected and expression value is above the average expression across all identities.

**Fig. S2: Differential gene usage across lifecycle. A**) Dotplot of the set of differentially expressed genes across life cycle stages. **B**) TOP go evaluation of all differentially expressed genes across life cycle stages. See supplementary Data S1.2 for full gene list

**Fig. S3: Full cell state complement**. **A**) UMAP colored by sample identity. **B**) UMAP representation of all cluster identities, separated by life cycle stage.

**Fig. S4: Digestive gland population**. **A**) Full dataset UMAP showing position of cluster subset **B**) UMAP distribution of identified clusters. **C**) Bar plot showing distribution of digestive gland cells across life cycle stages and medusa tissue samples as absolute cell numbers. **D**) Gene expression dot plot showing expression profile of *ashC* specifically in identified digestive gland cell subtypes. **E**) Gene expression dot plot showing expression profile of top 10 differentially expressed genes for each cluster, separated by life cycle contribution. Full gene list is available in supplementary data S1.10. **F**) *in situ* hybridization showing localization of chitinase gene expression to the gastric filaments

**Fig. S5: Mucin gland population**. **A**) Full dataset UMAP showing position of cluster subset **B**) UMAP distribution of identified clusters. **C**) Bar plot showing distribution of mucin gland cells across life cycle stages and medusa tissue samples as absolute cell numbers. **D**) Gene expression dot plot showing expression profile of *ngn* specifically in identified mucin gland cell subtypes. One paralog of the atoh8 gene family is also expressed specifically in mucin-subtypes. **E**) Gene expression dot plot showing expression profile of top 10 differentially expressed genes for each cluster, separated by life cycle contribution. Full gene list is available in supplementary data S1.7

**Figure S6: Cnidarian expansion of myc-family members. A)** Gene tree: Aurelia is indicated in blue, *Nematostella* in salmon, other cnidarians in black and bilaterian orthologs in grey. Expression within the two shared cell type states (early and mature) as indicated in Figure 3 are plotted. Internal nodes indicate bootstrap values. Ac: *Aurelia coerulea*; Bf: *Branchiostoma floridae*; Ce: *Caenorhabditis elegans*; Cx: *Cassiopea xamachana*; Ed: *Exaiptasia diaphana*; Hs: *Homo sapiens*; Hv: *Hydra vulgaris*; Hy: *Hydractinia symbiolongicarpus*; Mg: *Magallana gigas*; Nv: *Nematostella vectensis*; Of: *Owenia fusiformis*; Re: *Rhopilema esculentum*; Sm: *Sanderia malayensis*; Sp: *Stylophora pistillata*; Tc: *Tripedalia cystophora* **B**) Expression of included gene models on the *Hydra vulgaris* dataset (from https://research.nhgri.nih.gov/HydraAEP/SingleCellBrowser/).

**Fig. S7: Over-represented biological processes in differential gene lists (topGO)**. Each set of up-regulated genes from all neuroglandular subtypes was evaluated for over-represented biological processes with the topGO package. Full gene lists are available in supplementary data S1.8.

**Fig. S8: Expression of *atoh8* family transcription factors. A**) Expression profiles across all identified cell type families of each member of the *atoh8* family. **B:D**) Expression profiles of *atoh8* family genes within the neural clusters of *Aurelia* (B), *Nematostella* (C), and *Hydra* (D) highlights both broad expression profiles, as well as cell type specific expression. **E**) Gene family tree with *Nematostella* (salmon), *Aurelia* (blue) and human (red) paralogs highlighted. Internal nodes indicate bootstrap values. Ac: *Aurelia coerulea;* Bf: *Branchiostoma floridae*; Ce: *Caenorhabditis elegans*; Cx: *Cassiopea xamachana*; Ed: *Exaiptasia diaphana*; Hs: *Homo sapiens*; Hv: *Hydra vulgaris*; Hy: *Hydractinia symbiolongicarpus*; Mg: *Magallana gigas*; Nv: *Nematostella vectensis*; Of: *Owenia fusiformis*; Re: *Rhopilema esculentum*; Sm: *Sanderia malayensis*; Sp: *Stylophora pistillata*; Tc: *Tripedalia cystophora*

**Fig. S9: Muscle-specific transcription factors.** Dotplot of expression of the top differentially expressed gene models with DNA binding sites specific to the striated muscle subset. Full gene list is available in supplementary Data S1.4b

### Tables

#### Data S1: Differentially upregulated genes for all cell clusters

1. Medusa-specific genes from Figure 1C b) filtered list of Medusa-specific genes illustrated in Figure 1E.
2. Life cycle, all genes, b) DNA binding domain genes (putative transcription factors: pTF).
3. Coarse clustering (Figure 2A) b) pTF
4. Striated Muscle b) pTF
5. Epidermis b) pTF
6. Gastrodermis b) pTF
7. Mucin gland b) pTF
8. Neuroglandular b) pTF
9. Cnidocytes b) pTF
10. Digestive glands b) pTF
11. Muscle-specific gene sets

#### Data S2: Relevant technical information

1. Sequence mapping statistics
2. Gene model annotations
3. Coarse clustering
4. Epidermis
5. Gastrodermis
6. Digestive glands
7. Mucin gland
8. Cnidocytes
9. All.Neuroglandular
10. Striated Muscle
11. Aurelia vs Nematostella 1:1 OMA orthologs
12. In situ primer sequences

#### Data S3: Multiple sequence alignment: bHLH family of proteins

## Notes

### Competing Interest Statement

The authors have declared no competing interest.

### Summary of Updates

We have updated the mapping of the single cell data to a chromosome-level genome assembly of A. coerulea (NCBI). Additionally we have updated the gene models and consolidated these with our previous gene set to arrive at an improved gene annotation set. This includes the hand-curated gene annotation of about 5000 previously unannotated gene models. We subsequently re-ran all analyses on the single cell dataset. While the main messages of the previous version largely remained unchanged, we would like to stress that the new mapping tool resulted in a reduction of transcript models and overall improved data quality in our analyses.

