## Supplementary Figures HTML for "The emergence of medusa-specific cell states in the scyphozoan *Aurelia coerulea*": Supplement.html

Aurelia Atlas | Supplementary Data Figures


### Aurelia Atlas | Supplementary Data Figures

###### AlisonGCole

**Aurelia Cell Atlas:**

Supplementary materials in support of the conclusions drawn in the
manuscript.

**Gene duplications facilitated the emergence of
medusa-specific cell states in the scyphozoan Aurelia coerulea**
  
  
 Oliver Link1, Stefan M. Jahnel2, Kristin Janicek1, Daniel
Guerguerian1, Johanna Kraus1, Juan Daniel Montenegro1, Bob Zimmerman1,
Brittney Wick3, Konstantin Khalturin4, Alison G. Cole1, and Ulrich
Technau1,5

##### **Figure S1: Life cycle: medusa genes**

Figure S1: **A**) Total number of genes (nFeatures\_RNA)
and **B**) molecules (nCounts\_RNA) recovered from all cells
captured across life cycle stages **C**) Distribution of
medusa-specific genes across all cell types. The set of all
medusa-specific genes were considered. For every cluster the gene is
counted as expressed if reads are detected and expression value is above
the average expression across all identities.

##### **Figure S2: Life cycle: differentially expressed genes**

**Fig. S2: Differential gene usage across lifecycle.
A**) Dotplot of the set of differentially expressed genes across
life cycle stages. **B:E**) TOP go evaluation of all
differentially expressed genes across life cycle stages. See
supplementary Data S1.2 for full gene list

##### **Figure S3: Full cell state complement**

**Fig. S3: Full cell state complement**. UMAP
representation of all cluster identities, separated by life cycle
stage.

##### **Figure S4: Digestive Gland**

**Figure S4: Digestive gland population**.
**A**) Full dataset UMAP showing position of cluster subset
**B**) UMAP distribution of identified clusters.
**C**) Bar plot showing distribution of digestive gland
cells across life cycle stages and medusa tissue samples as absolute
cell numbers. **D**) Gene expression dot plot showing
expression profile of *ashC* specifically in identified digestive
gland cell subtypes. **E**) Gene expression dot plot
showing expression profile of top 10 differentially expressed genes for
each cluster, separated by life cycle contribution. Full gene list is
available in supplementary data S1.10 **F)** *in
situ* hybridization showing localization of chitinase gene
expression to the gastric filaments.

##### **Figure S5: Mucin gland cluster**

**Figure S5: Mucin gland population**.
**A**) Full dataset UMAP showing position of cluster subset
**B**) UMAP distribution of identified clusters.
**C**) Bar plot showing distribution of mucin gland cells
across life cycle stages and medusa tissue samples as absolute cell
numbers. **D**) Gene expression dot plot showing expression
profile of mucin-like-protein (*mlp-like*) and the bHLH
transcription factor *ngn* specifically in identified mucin gland
cell subtypes. One paralog of the *atoh8* gene family is also
expressed specifically in mucin subtypes *mucin.3* and
*mucin.medusa*.

#### **Figure S6: Myc gene tree**

**Figure S6: Cnidarian expansion of myc-family members.
A**) Maximum likelihood tree; *Aurelia* is indicated in
blue, *Nematostella* in salmon, other cnidarians in black and
bilaterian orthologs in grey. Expression within the two shared cell type
states (early and mature) as indicated in Figure 3 are plotted. Internal
nodes indicate bootstrap values. Ac: *Aurelia coerulea*; Bf:
*Branchiostoma floridae*; Ce: *Caenorhabditis elegans*;
Cx: *Cassiopea xamachana*; Ed: *Exaiptasia diaphana*; Hs:
*Homo sapiens*; Hv: *Hydra vulgaris*; Hy: *Hydractinia
symbiolongicarpus*; Mg: *Magallana gigas*; Nv:
*Nematostella vectensis*; Of: *Owenia fusiformis*; Re:
*Rhopilema esculentum*; Sm: *Sanderia malayensis*; Sp:
*Stylophora pistillata* ; Tc: *Tripedalia cystophora*
**B**) Expression of included gene models on the *Hydra
vulgaris* dataset (from *https://research.nhgri.nih.gov/HydraAEP/SingleCellBrowser/*).

##### **Figure S7: Neuron GO terms**

**Figure S7: Over-represented biological processes in
differential gene lists (topGO)**. Each set of up-regulated genes
from all neuroglandular subtypes was evaluated for over-represented
biological processes with the topGO package. Full gene lists are
available in supplementary data S1.8.

##### **Figure S8: ATOH8 family genes**

**Figure. S8**: Expression of atoh8 family transcription
factors. **A**) Expression profiles across all identified
cell type families of each member of the atoh8 family.
**B:D**) Expression profiles of atoh8 family genes within
the neural clusters of *Aurelia* (B), *Nematostella* (C),
and *Hydra* (D) highlights both broad expression profiles, as
well as cell type specific expression. G023106 is the hydra
*pou4* ortholog indicating class n2-type neurons, and G017257 is
the *ins* ortholog, indicating class n1-type neurons.
**E**) Gene family tree with *Nematostella*
(salmon), *Aurelia* (blue), *Hydra* (blue-green) and human
(red) paralogs highlighted. Internal nodes indicate bootstrap values.
Ac: *Aurelia coerulea*; Bf: *Branchiostoma floridae*; Ce:
*Caenorhabditis elegans*; Cx: *Cassiopea xamachana*; Ed:
*Exaiptasia diaphana*; Hs: *Homo sapiens*; Hv: *Hydra
vulgaris*; Hy: *Hydractinia symbiolongicarpus*; Mg:
*Magallana gigas*; Nv: *Nematostella vectensis*; Of:
*Owenia fusiformis*; Re: *Rhopilema esculentum*; Sm:
*Sanderia malayensis*; Sp: *Stylophora pistillata* ; Tc:
*Tripedalia cystophora*

##### **Figure S9: Muscle Transcription Factors**

**Figure S9:** **Muscle-specific transcription
factors.** Dotplot of expression of the top differentially
expressed gene models with DNA binding sites specific to the striated
muscle subset. Full gene list is available in supplementary Data
S1.4b.
